# Forgetting Dynamics For Items of Different Categories

**DOI:** 10.1101/2022.11.14.516480

**Authors:** Antonios Georgiou, Mikhail Katkov, Misha Tsodyks

## Abstract

How the dynamic evolution of forgetting changes for different materials is unexplored. By using a common experimental paradigm with stimuli of different types, we were able to directly cross-examine the emerging dynamics and we found that even though the presentation sets differ minimally by design, the obtained curves appear to fall on a discrete spectrum. Furthermore, we have previously proposed a model of forgetting based on the notion of retrograde interference with a single integer parameter. All measured curves were compatible with the model with different values of the parameter, hinting to a potential common underlying mechanism of forgetting.

---

Ever since Ebbinghaus’ seminal work (Ebbinghaus [1964]), quantitative measurements of performance are the staple of the studies of human memory (see e.g. Kahana [2012]). This endeavor is hindered by sensitivity of memory to multiple factors such as material being presented to participants, presentation protocols, age of the participants etc., that seem to preclude any universal characterization of performance. Following Dudai [2004], we consider memory as a 3-staged process consisting of (i) acquisition of information and its encoding, (ii) retainment it in memory over time and finally (iii) recall (we can schematically describe it as *A* → *M* → *Rec*). In the following, we assume for simplicity that each of the process results in a binary result, for example each presented item is either encoded in long-term memory or not. If a participant is presented with a list of *L* words, some number of them are either missed or not efficiently encoded (i.e. *A < L*), those encoded can be erased during the presentation of subsequent words (*M < A*) and finally some number of retained words are recalled (*Rec* < *M*). All three processes could contribute to the unpredictability of memory; however, in our recent publication (Naim et al. [2020]) we found that if we can experimentally estimate *M* for a group of participants, their average recall performance can be predicted surprisingly well by a simple phenomenological model of recall, resulting in the universal (i.e. independent on the experimental conditions and type of material presented) formula that relates *Rec* to *M* :

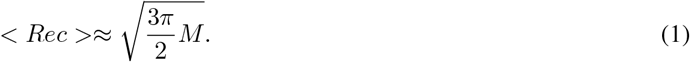

This implies that under manipulations performed in (Naim et al. [2020]) (different number of presented items, presentation speed and item category), most of the sensitivity of recall to experimental conditions is explained by their effect on *M*, i.e. on memory acquisition and maintenance during the presentation of the material to be remembered. Indeed the retention of a presented item in memory requires that (i) it is effectively acquired and stored in memory upon presentation and (ii) is not erased due to interference with subsequent acquisition of new items (forgotten; see Wixted [2004]). In the current study, we focus on the dynamics of forgetting when participants are exposed to a stream of items of a particular type. Following Ebbinghaus, we study the retention curve (denoted *R*(*t*) below), which, mathematically, describes the probability that an acquired item still remains in memory after time *t*. Since memory and forgetting are processes that happen in time, exploring their dynamics is of critical importance for probing putative substrate mechanisms of forgetting. In particular, we aim to establish how universal the forgetting dynamics is for different classes of inputs and different presentation conditions.

On the experimental front, even though there have been many studies that explore memory differences between different categories of stimuli such as words, pictures and sentences (e.g Jenkins et al. [1967], Nelson et al. [1976], Shepard [1967]), within category differences such as manipulating the level of abstraction within pictures (e.g Bellhouse-King and Standing [2007], Goldstein and Chance [1971]), the conceptual and schematic similarity (Nelson et al. [1976]) or the distinctiveness of the material in the format it is presented (Ensor et al. [2019]), in most cases memory performance has been addressed as a singular point in time. Rigorous testing of any mathematical model of forgetting as a process though, requires diverse data as a function of time in accordance with the retention function, which in turn provide a stricter ‘fitness’ criterion. An important question not addressed in previous studies is how generic the shape of the forgetting curve is for different material. We previously demonstrated some degree of universality for the performance of *recall* by measuring the relation between the number of remembered versus recalled items, acquired in experiments with random lists of words or short sentences Naim et al. [2020]. Could the retention curve exhibit the same universality across stimuli types? The results mentioned above argue against this, but universality could still be observed if the retention curves for different stimulus types are scaled versions of each other, i.e. have similar shapes and only differ in absolute values.

Comparison between forgetting curves has been a contentious subject however, starting with the debate between Slamecka (Slamecka and McElree [1983]) and Loftus (Loftus [1985]) on whether initial learning affects the rate of forgetting with Wixted Wixted [1990] trying to resolve it by posing the issue as a disagreement of definition of the forgetting rate. To avoid any confusion, we consider forgetting as the change in performance, namely the proportion of correct responses in recognition tasks, as a function of the number of interposed items between presentation and test (lag). We do not examine differences in physical time, but rather we consider a time unit as the distance between two events. In principle we follow Standing’s single-trial tradition (Standing [1973]) where he investigates the relationship between items in memory versus items presented, although we test for the full spectrum of the presentation range rather than at predefined points. Moreover, we do not attempt to fit the data to a candidate retention function. Instead, we focus on whether psychological principles can lead to mathematically generated curves that are compatible with experimental observation.

Theoretical work has suggested a variety of candidate mechanisms of forgetting with accompanying mathematical models, such as the passive decay of memories (e.g Kahana and Adler [2017]), temporal distinctiveness (e.g Brown and Lewandowsky [2010]) and interference (e.g Georgiou et al. [2021]) to name a few. (For an extensive review of decay and interference literature, see Wixted [2004]). These theoretical results should then be juxtaposed and compared with experimental findings to assess the validity of their claims. In particular, memory performance varies with the material used in each experiment (see below) and theoretical models should reflect that.

We recently introduced a mathematical model of forgetting based on retroactive interference between acquired memories that is hypothesized to depend on their relative ‘valence’ measures. More precisely, each memory is categorized with a multidimensional valence and a newly acquired memory erases all the previous ones that has lower valence in all dimensions Georgiou et al. [2021]. The inspiration behind having memory items with valences in multiple independent dimensions comes from the observation that at the encoding level a single item can elicit responses in multiple contextual clusters (Huth et al. [2016]). Similarly at the retrieval level, engram complexes representing a single memory have been reported to be distributed across cortical regions (Roy et al. [2022]). If all the valences are assumed to be randomly sampled from an arbitrary distribution, this model can be analytically solved for the retention function. We compared the model predictions with the results of experiments with presentation of streams of randomly chosen common nouns interrupted for recognition trials. The experimentally obtained retention function was shown to be similar to that of the 5-dimensional version of our model. As opposed to the recall model of Naim et al. [2020] that had no free parameters, the forgetting model has one parameter that is the dimensionality of the memory valences, and hence could potentially account for different shapes of retention function. In the current contribution, we repeated the experiments of Georgiou et al. [2021] with other types of materials, both verbal and visual, to access the diversity of retention functions across stimulus types and the generality of the retrograde forgetting model of Georgiou et al. [2021].

The model of forgetting proposed in (Georgiou et al. [2021]) characterizes each presented item as an n-dimensional vector, with components sampled from an arbitrary n-dimensional distribution. The individual components of this vector represent the valence, or more simply the memory strength of each particular item in different dimensions. We assume that items are acquired at each time step and committed to memory. Every newly acquired item interacts retroactively with already stored items and erases all of those that are weaker than it element-wise in each dimension (see Figure 1). Under the simplifying assumption that all components of memory valences are sampled independently from each other, the probability that an item is retained in memory t time steps after its acquisition, i.e. the value of the retention function *R*_*n*_(*t*), can be expressed iteratively as:

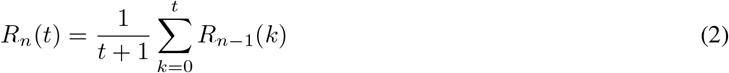

where n is the number of dimensions and *R*_1_(*t*), the retention curve of the 1-dimensional model is simply:

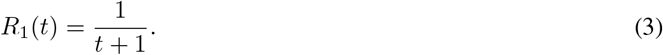

Equation 3 is trivially obtained by noting that for *n* = 1, the probability that none of the *t* next items will erase the currently acquired one is the same as the chance that out of *t* + 1 independent random numbers sampled from the same probability distribution, the first one is the largest. This chance is clearly given by 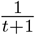 since each sample can be the largest one with the same probability. The recursive equation 2 can be obtained by noting that if we only consider the first dimension, then out of next *t* presented items any number of them (*k*) from 0 to *t* can be larger than the present item with equal probability, which is therefore given by 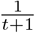. On the other hand, if there are exactly *k* such items, only they can potentially erase the current one (the rest have smaller valence along the first dimension). The current items can therefore only survive the next *t* steps due to other *n* − 1 dimensions, and the probability for this is then by definition given by *R*_*n*−1_(*k*). Averaging over all possible values of *k* results in equation 2.

**Figure 1:**
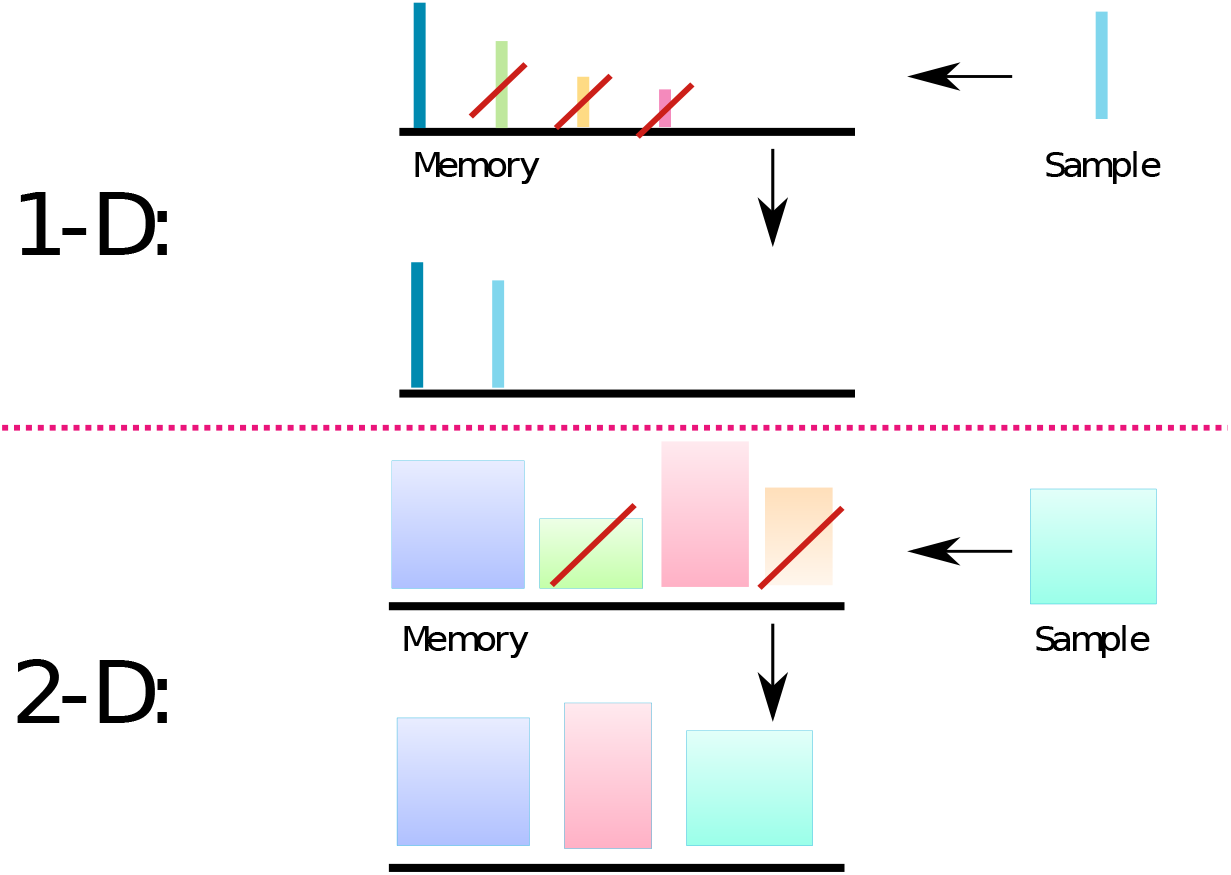
Interference model of forgetting. **1-D**. Each item is represented as a thin vertical bar. The height of the bar corresponds to the valence of an item. The top row bars above the black line represent items that are stored in memory just before the acquisition of a new item, shown on the right (Sample). All the items that have smaller valence (bar height) than the new item are discarded from memory (crossed by red bar), and the new item is added. Bottom row represents the memory content, after the new memory is acquired. **2-D**. Same as 1-D, but each memory item has 2 valences, represented by the width and height of a rectangular. In this case, all the items that have both valences smaller than the corresponding valences of the new item are discarded.

An equivalent closed-form solution to the retention function can also be found to be

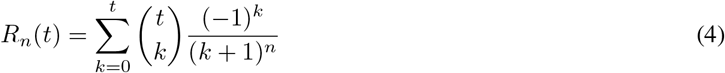

where 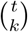 is binomial coefficient (Katkov et al. [2022]). The model yields a family of distinct curves by varying the free integer parameter *n*, with higher *n* leading to better retention, i.e. higher values of retention function (see Figure 3 below). Recognition experiments’ data for a list of nouns were well described by the model with *n* = 5 Georgiou et al. [2021].

We expanded the investigation of Georgiou et al. [2021] by conducting similar recognition experiments while changing the presentation material to examine the differences between the retention curves. Participants were recruited online in Amazon’s Mechanical Turk® platform and were requested to attend to a stream of 200 items at a rate of 1.5s per item. At random moments throughout, presentation was paused and a two alternative forced choice (2AFC) recognition task was introduced. Participants were given a choice of two items, one that had appeared before and another one which was not previously shown (see Figure 2). They were instructed to select the one they remembered as previously seeing. The targets of the 2AFC queries were of two kinds. One kind was comprised of the first ten presented items. These items were inquired at random order and at random times throughout the experiment yielding ten responses per participant to 2AFC tasks with a lag spanning up to 199 items. The results of this kind query are depicted in Figure 3 where the probability of a correct response (*C*) is plotted versus the number of intervening items between presentation and test. As in Georgiou et al. [2021] and in many previous studies, we related the probability of correct response after a certain number *t* of intervening presentations to the probability that the corresponding item is still in memory (i.e. retention function *R*(*t*)) by the following equation:

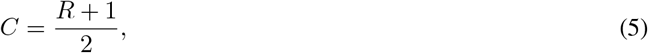

which follows from the assumption that if the item is in memory, a participant chooses the correct target at the recognition test; otherwise, the participant chooses an item randomly. The other kind of recognition query was used to select participants who were attentive for the whole duration of the experiment. Another ten 2AFC queries were presented at random times during the experiment, only in this case the inquired item was always the one before last item (2-back task). This item should still be in working memory and attentive participants were expected to have 100% accuracy in these tasks. Only the participants that achieved a perfect score for the 2-back task were included in the analysis.

**Figure 2:**
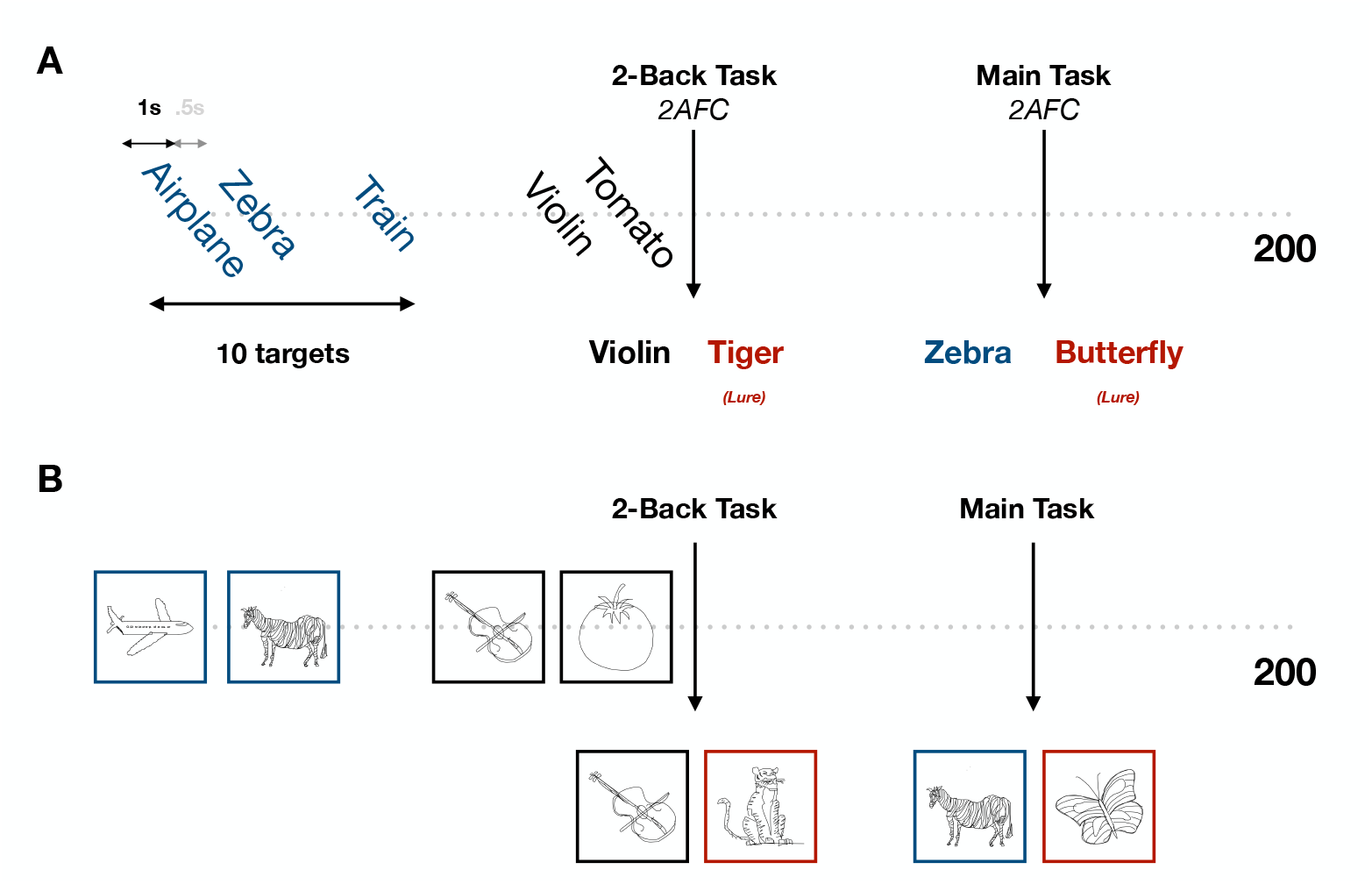
Experimental Protocol: Items are presented sequentially for 1s followed by a 0.5s of blank screen. At random points the presentation is paused and a two-alternative forced choice recognition task is shown. The target (stimulus previously shown) comes from one of the first 10 presented items (Main Task), or the second to last shown item (2-Back Task) which is used to filter inattentive participants. The distractor (stimulus not previously shown, or ‘lure’) is shown in red. Stimuli examples of the nouns experiment are shown in **A** and of the sketches one in **B**.

**Figure 3:**
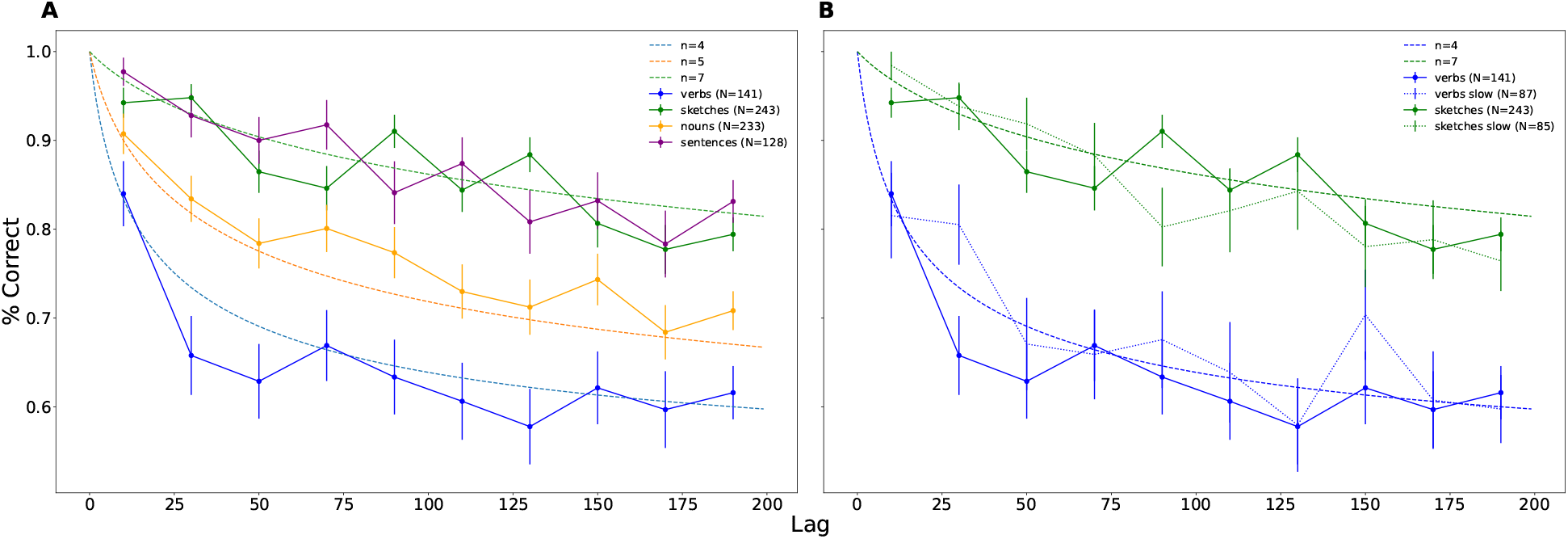
Experimental Retention Curves For Lists of Different Types And Corresponding Theoretical Curves With Different Parameter Values. **A:** Experimental curves produced from a **2AFC** memory task on lists of 200 items for different types of material (Blue: Verbs, Orange: Nouns, Purple: Sentences, Green: Sketches) with a presentation speed of 1s (where the stimulus is shown) plus 0.5s inter-item interval (blank screen). The dashed lines correspond to theoretical curves produced by our retroactive interference model for different values of the dimensionality parameter *n* (Purple: *n* = 4, Brown: *n* = 5, Pink: *n* = 7). **B:** Similar to **A** for verbs and sketches with 2 different presentation speeds. The solid lines represents the results for verbs and sketches as they appear in **A**, while the dotted lines represent the same experiment as in **A** but with a presentation speed of 2+0.5s. Again, the dashed lines depict the model’s theoretical curves. The data produced from a longer presentation time, do not differ substantially from their faster counterparts. In both **A** and **B**, the number of participants contributing to each curve is denoted by the value of N in the bracket next to each category.

To examine the difference between verbal and pictorial stimuli we conducted identical experiments using the same items, only in one case they appeared as sketches and in the other as words (labels; see Figure 2 for some examples). The sketches were simple black and white human-drawn line drawings of different objects while the names of these objects were used in the word experiment. The resulting retention curves of these two experiments can be seen in Figure 3 **A** where the orange curve represents the sketch condition and the blue represents the word one. The two conditions give rise to two distinct non-overlapping curves, with the performance for sketches being consistently higher for every time bin. Comparing these two curves with the curves generated from our model, we see that the results for sketches follow reasonably well the theoretical curve with a dimension of 7, while the words lie close to the one with the dimension of 5 (even though admittedly above it). In our previous study Georgiou et al. [2021] where we also used nouns, retention curve was closer to the theoretical curve of dimension 5 then in the current study. We speculate that the difference in results could be due to different pools of nouns used in the experiments, in particular in the current study we only used the concrete nouns that depict objects, and people could use visual imagery to better remember them.

A question that arises is whether the observed difference in retention of different stimulus categories is simply the product of verbal versus pictorial stimuli. To address this, the next experiment was conducted using trains of sentences as stimuli. These sentences were small pseudo-definitional, matter-of-fact statements, pertaining to a single word such as ‘Consumers buy food’. The resulting curve can be observed again in Figure 3 **A** (green curve). The general performance is higher than the word condition as might have been expected, but more interestingly, the curve is almost indistinguishable from the sketch curve, lying around the theoretical curve for *n* = 7.

Following the same train of thought, we introduced another set of stimuli to the same experiment. In our words condition, the list was comprised of nouns as is typical of recognition experiments of this kind. A minimal modification to this experiment would be to substitute nouns to verbs, taking care to exclude verbs that double as nouns. Verbs were previously shown to be remembered worse than nouns when participants were exposed to natural material like phrases or passages (see e.g. Reynolds and Flagg [1976], James [1972], Wearing [1970]) but comparisons between lists of verbs and nouns were not performed to the best of our knowledge. Indeed we found a poorer recognition for verbs, with the resulting retention function quite close to the one predicted by our model with *n* = 4 (see Figure 3 **A**).

Our interpretation of the above results and their comparison to the theoretical model of Georgiou et al. [2021] depends crucially on the assumption that initial degree of encoding of presented items in long-term memory does not systematically differ for different stimulus categories. In other words, that initial recognition of items immediately after acquisition, if you disregard the working memory effects, is the same for all stimuli (in other words, extrapolation the retention curves to *t* = 0 is close to perfect independently on the type of the stimulus, for participants who pay a close attention to all the presented stimuli). An alternative interpretation of the results could be that e.g. verbs are encoded in long-term memory differently than other stimuli such that their immediate recognition is less precise, and this in turn results in faster rate of subsequent forgetting. We speculated that in this case, if one increase the presentation time of verbs, the initial encoding will be stronger and the retention curve will move in the direction of other stimuli. We therefore performed additional experiments where we increased the presentation time for each verb from 1 to 2 seconds. Our results showed however that while the relative number of participants with perfect 2-back recognition went down (possibly because of the longer overall time of the experiment), their retention curves did not change significantly except for one point (see Figure 3 **B**). For the sake of generality, we performed the same manipulation with sketches and again obtained very similar retention curves for both presentation speeds. We believe that our results point to quite a high degree of universality in forgetting dynamics, namely when controlled for acquisition (in our case, only considering participants who attend to each stimulus) and stimulus category, retention curves exhibit generic shape that does not seem to depend on other experimental conditions. It remains to be seen whether this conclusion will hold for other experimental manipulations not considered in this study, such as the age of participants.

Taken together with the results of Naim et al. [2020], we conclude that two of the processes involved in memory for random lists of items, namely retainment and recall, exhibit quite a significant degree of universality. The dependency of retention dynamics on the type of stimuli presented for remembering contrasts retention with recall that appears to be independent on the item category. One possible explanation for category-dependent retention is that different stimulus types are processed by different number of brain areas (see e.g. Hasson et al. [2002]). We also note that both these models are deterministic, e.g. given the exact values for each presented stimuli, we could predict exactly which one of them will be erased and at which time. How close is this to reality is an open interesting question, as well as whether extending the model to allow for probabilistic interference, possibly depending on the difference in valences between the items could provide a better account of the data.

Retention curves could reveal important insights into mechanisms of forgetting and provide a testing ground for theoretical models. In the present study, we have used the recognition experimental paradigm to examine different, but not unrelated, types of material. Interestingly, we see that within the verbal domain, with a difference as minimal as nouns versus verbs, two distinct curves emerge. On the other hand, even between domains, in our case verbal (sentences) and pictorial (sketches), the resulting curves are practically indistinguishable. An interesting overarching observation on all the stimulus categories that we considered in this study is that the retention curves appear to be either quite distinct or fully overlapping. Interestingly, this observation is consistent with our model of forgetting that has one discrete parameter, namely the number of dimensions of the memory valence. Moreover, all the data show compatibility with the model for different number of dimensions, from 4 for lists of verbs up to 7 for lists of sketches and sentences. We are therefore tempted to speculate that possible retention curves for all types of stimuli form a discrete set of ‘universality classes’ rather than a continuum. More experiments with different categories of items should be performed to either confirm or reject this speculation.

In each experiment conducted in this study, we have looked on retention curves that emerge from presented material coming strictly from a single category. Every presentation train contained only sketches or only nouns for example. In future studies, there are many categories to explore apart from the narrow selection considered here, such as sounds and natural scenes. Outside of experimental conditions, however, people are bombarded with input comprised of different categories, in different levels of abstraction and in different modalities, simultaneously and sequentially. It would be reasonable to expect that this diverse information would interact and interfere with each other. However, assuming the forms of the resulting retention curves is a non-trivial task. It would be interesting to examine whether these new ‘complex’ curves remain a discrete set or populate the entire retention space.

## Materials and Methods

All experiments were conducted through crowd-sourcing, utilising Amazon’s Mechanical Turk®platform. Ethics approval was obtained by the Institutional Review Board of the Weizmann Institute of Science and each participant accepted an informed consent form before participation. The experiments had a duration of approximately six minutes and a compensation of one dollar was rewarded upon completion of the task. The format of the experiment was identical for every stimulus set. Participants were greeted with an instruction page which gave a brief explanation of the task at hand. Upon pressing the start button, participants were presented with a train of 200 randomly selected items from a larger corpus, shown centrally with a white background, at a rate of 1.5s per item, with the item being visible for 1s plus 0.5s of blank screen between items. The first ten presented items were considered the target items. Throughout the presentation segment and at random times the presentation of items was paused until a two alternative forced choice task was completed. Two items were presented with the instruction to choose the one that the participant remembered seeing before. One of the two items belonged to one of the targets, i.e one of the first ten presented items, and the other was a lure, an item not included in the presentation list but from the same original set. The position of the correct item versus the lure was chosen at random at each given instance. The target items were probed at a random order and not in the order they were presented. All recognition tasks appeared at random times during the presentation of the items apart from the last one which always occurred at the end of the presentation. In order to discard inattentive participants, apart from the ten recognition trials pertaining to the targets another set of ten trials was introduced, inquiring about the item presented previous to the last one versus a lure. This item should still be in working memory and we would expect a perfect score from fully engaged participants. These trials were also spread throughout presentation at random times.

### Verbs

The list of verbs was generated by selecting all the verbs from the WordNet database. The verbs that double as nouns were excluded, like for example smoke. Then they were ordered according to frequency based on a variety of online corpora and the 1000 most frequent were selected.

### Sketches And Labels

The list of sketches was curated by hand from the database of human drawings created in Eitz et al. [2012]. A single sketch was selected from each category and the corresponding category name was used to comprise the list of labels.

### Sentences

The sentences were manually created based on the list of words used in Georgiou et al. [2021]. From each word, a small pseudo-definitional sentence was created with the purpose to not convey further information than the word in solitude would. One example of such a sentence was ‘BOATS TRAVEL ON WATER’ created from the word ‘BOAT’. The fact that a boat travels on water is implicit information already contained in the word itself.

### Analysis

Analysis was conducted through custom Python scripts. The lag was computed as the difference between a query position and a presentation position in the presentation stream. The results were then binned in time (lag) to generate mean values of correct recognition per bin (10 bins in total) which gave rise to the retention curves. Not all participants had queries for all bins. To account for guessing in the recognition tasks we assumed that if a person has forgotten the inquired word then she/he would have to randomly select one of the two options with equal probability, while if she/he remembered that item, then the correct option would be chosen. Therefore in order to account for this guessing, we express recognition performance as:

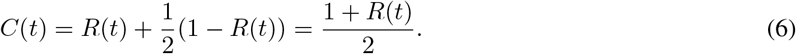

## Acknowledgements

This research has received funding from the European Union’s Horizon 2020 Framework Programme for Research and Innovation under the Specific Grant Agreement No. 785907 (Human Brain Project SGA2); EU-M-GATE 765549 and Foundation Adelis. We thank Dr. Rafi Malach for helpful discussions.

## References

Hermann Ebbinghaus. Memory: A contribution to experimental psychology. Dover, New York, 1964.

Michael Jacob Kahana. Foundations of human memory. OUP USA, 2012.

Yadin Dudai. The neurobiology of consolidation, or, how stable is the engram? Annual Review of Psychology, 55(4): 51–86, 2004.

Michelangelo Naim, Mikhail Katkov, Sandro Romani, and Misha Tsodyks. Fundamental law of memory recall. Physical review letters, 124(1):018101, 2020.

John T Wixted. The psychology and neuroscience of forgetting. Annu. Rev. Psychol, 55:235–69, 2004. doi:10.1146/annurev.psych.55.090902.141555. URL http://www.annualreviews.org.

Joseph R Jenkins, Daniel C Neale, and Stanley L Deno. Differential memory for picture and word stimuli. Journal of Educational Psychology, 58(5):303, 1967.

Douglas L Nelson, Valerie S Reed, and John R Walling. Pictorial superiority effect. Journal of experimental psychology: Human learning and memory, 2(5):523, 1976.

Roger N Shepard. Recognition memory for words, sentences, and pictures. Journal of verbal Learning and verbal Behavior, 6(1):156–163, 1967.

Mathew W Bellhouse-King and Lionel G Standing. Recognition memory for concrete, regular abstract, and diverse abstract pictures. Perceptual and motor skills, 104(3):758–762, 2007.

Alvin G Goldstein and June E Chance. Visual recognition memory for complex configurations. Perception & Psychophysics, 9(2):237–241, 1971.

Tyler M Ensor, Aimée M Surprenant, and Ian Neath. Increasing word distinctiveness eliminates the picture superiority effect in recognition: Evidence for the physical-distinctiveness account. Memory & cognition, 47(1):182–193, 2019.

Norman J Slamecka and Brian McElree. Normal forgetting of verbal lists as a function of their degree of learning. Journal of Experimental Psychology: Learning, Memory, and Cognition, 9(3):384, 1983.

Geoffrey R Loftus. Evaluating forgetting curves. Journal of Experimental Psychology: Learning, Memory, and Cognition, 11(2):397, 1985.

John T Wixted. Analyzing the empirical course of forgetting. Journal of Experimental Psychology: Learning, Memory, and Cognition, 16(5):927, 1990.

Lionel Standing. Learning 10000 pictures. The Quarterly journal of experimental psychology, 25(2):207–222, 1973.

Michael J. Kahana and Mark Adler. Note on the power law of forgetting. bioRxiv., 2017. doi:10.1101/173765. URL https://www.biorxiv.org/content/early/2017/08/09/173765.

Gordon DA Brown and Stephan Lewandowsky. Forgetting in memory models: Arguments against trace decay and consolidation failure. In Forgetting, pages 63–90. Psychology Press, 2010.

Antonios Georgiou, Mikhail Katkov, and Misha Tsodyks. Retroactive interference model of forgetting. The Journal of Mathematical Neuroscience, 11(1):1–15, 2021.

Alexander G Huth, Wendy A De Heer, Thomas L Griffiths, Frédéric E Theunissen, and Jack L Gallant. Natural speech reveals the semantic maps that tile human cerebral cortex. Nature, 532(7600):453–458, 2016.

Dheeraj S Roy, Young-Gyun Park, Minyoung E Kim, Ying Zhang, Sachie K Ogawa, Nicholas DiNapoli, Xinyi Gu, Jae H Cho, Heejin Choi, Lee Kamentsky, et al. Brain-wide mapping reveals that engrams for a single memory are distributed across multiple brain regions. Nature communications, 13(1):1–16, 2022.

Mikhail Katkov, Michelangelo Naim, Antonios Georgiou, and Misha Tsodyks. Mathematical models of human memory. Journal of Mathematical Physics, 63(7):073303, 2022.

Allan G Reynolds and Paul W Flagg. Recognition memory for elements of sentences. Memory & Cognition, 4(4): 422–432, 1976.

Carlton T James. Theme and imagery in the recall of active and passive sentences. Journal of Verbal Learning and Verbal Behavior, 11(2):205–211, 1972.

AJ Wearing. The storage of complex sentences. Journal of Verbal Learning and Verbal Behavior, 9(1):21–29, 1970.

Uri Hasson, Ifat Levy, Mahrlene Behrmann, Talma Hendler, and Rafael Malach. Eccentricity bias as an organizing principle for human high-order object areas. Neuron, 34:479–490, 2002.

Mathias Eitz, James Hays, and Marc Alexa. How do humans sketch objects? ACM Trans. Graph. (Proc. SIGGRAPH), 31(4):44:1–44:10, 2012.

